# *luxA* gene from *Enhygromyxa salina* encodes a functional homodimeric luciferase

**DOI:** 10.1101/2023.02.27.530324

**Authors:** Anna Yudenko, Sergey V. Bazhenov, Vladimir A. Aleksenko, Ivan M. Goncharov, Oleg Semenov, Alina Remeeva, Vera V. Nazarenko, Elizaveta Kuznetsova, Vadim V. Fomin, Maria N. Konopleva, Rahaf Al Ebrahim, Nikolai N. Sluchanko, Yury Ryzhykau, Yury S. Semenov, Alexander Kuklin, Ilya V. Manukhov, Ivan Gushchin

## Abstract

Several clades of luminescent bacteria are known currently. They all contain similar *lux* operons, which include the genes *luxA* and *luxB* encoding a heterodimeric luciferase. The aldehyde oxygenation reaction is catalyzed by the subunit LuxA, while LuxB is inactive. Recently, genomic analysis identified a subset of bacterial species with rearranged *lux* operons lacking *luxB*. Here, we show that the product of the *luxA* gene from the reduced *luxACDE* operon of *Enhygromyxa salina* is luminescent upon addition of aldehydes both *in vivo* in *Escherichia coli* and *in vitro*. Overall, *Es*LuxA is less bright compared with luciferases from *Aliivibrio fischeri* (*Af*LuxAB) and *Photorhabdus luminescens* (*Pl*LuxAB), and most active with medium-chain C4-C9 aldehydes. Crystal structure of *Es*LuxA determined at the resolution of 2.71 Å reveals a classical monooxygenase fold, and the protein preferentially forms a dimer in solution. The mobile loop residues 264-293, which form a β-hairpin or a coil in *Vibrio harveyi* LuxA, form α-helices in *Es*LuxA. Phylogenetic analysis shows *Es*LuxA and related proteins may be bacterial protoluciferases that arose prior to duplication of the *luxA* gene and its speciation to *luxA* and *luxB* in the previously described luminescent bacteria. Our work paves the way for discovery of new luciferases that have an advantage of being encoded by a single gene.

## 1. Introduction

Luminescence is widespread in the living world, with diverse organisms using it for visual communication to scare off predators, attract prey, or in courtship behavior [1]. More than 30 types of bioluminescence systems, usually consisting of a protein called luciferase and its substrate, luciferin, are currently known [2], with new ones still being discovered [3] and being used for many applications, such as engineering biosensors and reporters [4,5] or developing autoluminescent plants [6].

One of the best studied bioluminescence systems is found in bacteria such as *Vibrio harveyi, Aliivibrio fischeri, Photobacterium mandapamensis, Photobacterium leiognathi, Photorhabdus luminescens* and others [7,8]. In the ecology of modern marine microbiota, bacterial bioluminescence is believed to be used for detoxification of reactive oxygen species [9], photoreactivation of DNA repair, providing cells with additional protection from the lethal effects of short-wave solar UV radiation [10], in symbiotic relationships with eukaryotic organisms [11] or for attracting them [12]. The active protein complex LuxAB is a heterodimer of α (LuxA) and β (LuxB) subunits. As a flavin-dependent monooxygenase [13], LuxAB binds reduced flavin mononucleotide (FMNH_2_) and utilizes molecular oxygen to convert a long chain aldehyde to a fatty acid and emit light. Alongside the *luxA* and *luxB* genes, corresponding operons also contain the genes encoding a NADPH-dependent acyl protein reductase (*luxC*), an acyl-transferase (*luxD*), and an acyl-protein synthetase (*luxE*), usually in the order *luxCDABE* [7,8]. LuxC and LuxE form a complex [14]. Additionally, *luxF* and *luxG* genes, encoding a myristylated flavin scavenger [15,16] and a flavin reductase, respectively, may also be present [7,8].

Overall, bacterial luciferases have been extensively studied. Crystallographic structure of *V. harveyi* LuxAB (*Vh*LuxAB) in complex with FMNH_2_ has been determined, revealing the general architecture of the heterodimer as well as the binding site of the flavin [17]. The structure also revealed presence of a long flexible loop covering the active site; this loop was shown later to be important for luminescence [18,19]. The reaction was shown to occur in the α subunit [17] starting with peroxide formation [20]. When expressed separately, the subunits display activity at the level of 10^-5^–10^-6^ of that of the heterodimer [21], and *V. harveyi* LuxA remains monomeric, whereas LuxB forms a homodimer [22]. Crystallographic studies of the latter revealed similar structure to that in the heterodimer [23,24]. Interestingly, *Photobacterium* LuxF also forms a homodimer with a similar monooxygenase fold [25–27]. LuxA and LuxB can be expressed in eukaryotic cells [28] and are active when fused [29].

Recently, a bioinformatic study revealed presence of luciferase operons in hundreds of genomes, and identified *luxA*-like genes in species that were not reported previously to display luminescence [30]. These *luxA*-like genes are located in putative *luxCEDA*, *luxCAED* and *luxAxxCE* operons that miss *luxB* [30]. Here, we describe a study of the product of the peculiar *luxA* gene from *Enhygromyxa salina*. We show that it is a functional homodimeric luciferase and describe the implications.

## 2. Materials and Methods

### 2.1. Bioinformatics

Seed alignment for the luciferase-like monooxygenase domain (Pfam family PF00296) was downloaded from Pfam [31] on April 12^th^, 2020. This alignment was used to build an initial PSSM matrix as an input for PSI-BLAST [32] to perform a search against the non-redundant set of sequences in the NCBI database (all non-redundant GenBank coding sequence translations, PDB, SwissProt, PIR and PRF excluding environmental samples from whole genome sequencing projects), also on April 12th, 2020. The cutoff E-value was chosen at 0.001. Resulted set of sequences was clustered using UCLUST [33] at the 40% identity level and centroids were used as a dataset for further phylogenetic analysis. Additionally, a set of known bacterial luciferase sequences was used to perform PSI-BLAST searches against the same database. Top 1000 results from this search were added into the dataset for further phylogenetic analysis. Finally, the resulting dataset was combined with the sequences representative of PF00296 family members with available experimentally determined structures. Multiple sequence alignment of the collected sequences was calculated using MAFFT v. 7.402 [34] with FFT-NS-2 algorithm [35]. Columns containing more than 50% gaps were removed using trimAl [36]. The phylogenetic tree was constructed using FastTree2 [37]. LG amino acid substitution model [38] was used, with 20 rate categories of CAT approximation, 44/22 rounds of ME/ML nearest-neighbor interchanges with turned off heuristics, 4 rounds of subtree-prune-regraft moves and the maximum length of a move of 10. Illustrations of the phylogenetic trees were prepared using FigTree v. 1.4.4 [39].

### 2.2. Bacterial strains and plasmids used in the study

Bacterial strains and plasmids used in the study are listed in Table 1. Detailed description of cloning for pSluxA, pABX, pABX-T7, and pLuxG-T7 plasmids is provided in supporting information.

**Table 1.** Bacterial strains and plasmids used in the study.

| Name | Description | Source |
| --- | --- | --- |
| Bacterial strains |  |  |
| *E. coli* K12 MG1655 | *F ilvG rfb-50 rph-1*;<br>was used to express the *luxAB* genes from *A. fischeri* and<br>*P. luminescens* for luminescence measurements | VKPM<br>(Russia) |
| *E. coli* BL21(DE3) | *F ompT hsdS<sub>B</sub> (r<sub>B</sub><sup>-</sup>, m<sub>B</sub><sup>-</sup>) gal dcm* (DE3);<br>was used to express the *luxA* gene from *E. salina*, *luxAB* from *P. luminescens*, *luxAB* from *A. fischeri*, and *luxG* from *V. aquamarinus*<br>for luminescence measurements | Novagen<br>(Germany) |
| *E. coli* C41(DE3) | BL21(DE3) phenotypically selected for tolerance to toxic proteins [67]; was used to overexpress the *luxA* gene from *E. salina* for structural studies | SigmaArdrich<br>(USA) |
| Plasmids |  |  |
| pABX | pUC19 with *luxAB* from *P. luminescens* [68] under promoter *Plac* (Ap<sup>r</sup>) | This study |
| pABX-T7 | pET15 with *luxAB* from *P. luminescens* under T7 promoter | This study |
| pLuxG-T7 | pET15 with *luxG* from *V. aquamarinus* VNB-15 (mentioned in [69])<br>under T7 promoter | This study |
| pF2 | pUC18 with *luxAB* from *A. fischeri* under promoter *Plac* (Ap<sup>r</sup>) | [70] |
| pSluxA | pET11 with *luxA* from *E. salina* under T7 promoter | This study |

### 2.3. Functional characterization in vivo

To characterize the luciferase activity, cell cultures were grown overnight on agarized (1.5% w/v) autoinduction ZYP-5025 medium [40] at 22 °С. Immediately prior to luminescence measurements, the overnight cultures were resuspended in liquid LB to reach the final optical density (OD) at 600 nm of 0.05 for *E. coli* K12 MG1655 pABX and K12 MG1655 pF2; and to OD of 2 for *E. coli* BL21(DE3) pSluxA, because of its very low luminescence. *E. coli* BL21(DE3) transformed with pET11 harboring a gene encoding a flavin-binding protein CagFbFP [41] was used as a negative control. The effect of aldehyde chain length on enzyme activity in investigated cells was obtained by subjecting the cell suspensions to different aldehydes (C1-C12 aldehydes were purchased from Sigma-Aldrich, USA; C13-C18 were purchased from Macklin, China) at the final concentration of 10 mM and then measuring the maximum light intensity using Biotox-7BM (BioPhysTech, Russia). Michaelis constants were determined by subjecting the cell suspensions to different aldehyde concentrations and fitting the luminescence curves using nonlinear least squares regression as implemented by Angel Herráez (https://biomodel.uah.es/en/metab/enzimas/MM-regresion.htm) based on the algorithm by John C. Pezzullo (https://statpages.info/nonlin.html).

### 2.4. Protein expression and purification

Isolation of luciferases *Es*LuxA and *Pl*LuxAB was performed using the same approach. The protein was expressed in *Escherichia coli* strain C41 (DE3). Cells were cultured in shaking baffled flasks in LB medium containing 150 mg/l ampicillin. Protein expression was induced with 1 mM IPTG and continued for 22 hours at 20 °C. Harvested cells were resuspended in phosphate-buffered saline with addition of 1 mM PMSF and disrupted in M-110P Lab Homogenizer (Microfluidics, USA) at 25000 psi. The lysate was clarified by removal of cell membrane fraction by ultracentrifugation at 40000 g for 1 h at 10 °C. The supernatant was incubated with Ni-nitrilotriacetic acid (Ni-NTA) resin (Qiagen, Germany) on a rocker for 1 hour at 4 °C. The Ni-NTA resin and supernatant were loaded on a gravity flow column and washed with buffer containing 300 mM NaCl and 50 mM Tris-HCl, pH 8.0, supplemented with 20 mM imidazole. The protein was eluted in a buffer containing 50-200 mM imidazole, 300 mM NaCl and 50 mM Tris-HCl, pH 8.0. The eluate was concentrated and subjected to size-exclusion chromatography (SEC) on Superdex® 200 Increase 10/300 GL column (GE Healthcare Life Sciences, USA) in a buffer containing 150 mM NaCl and 25 mM Tris-HCl, pH 8.0.

Isolation of LuxG was performed using the same approach with the following modifications. After cell lysis, LuxG was found predominantly in an aggregated state. Consequently, urea was added to the crude lysate to the final concentration of 2 M. The sample was centrifuged at 15000 g for 45 min prior to application of supernatant to Ni-NTA. The resulting LuxG protein had a concentration of about 0.2 mg/ml and exhibited enzymatic activity; it was further used to restore FMN in the luciferase reaction.

### 2.5. Functional characterization in vitro

*In vitro* measurements of luciferase activity of both *Es*LuxA and *Pl*LuxAB were carried out in 10 mM PBS buffer pH 7.4, supplemented with 20 μM FMN, 0.2 mM NADH, 0.4 μM LuxG, and aldehyde. To measure the activity of *Pl*LuxAB, decanal was added to the buffer at the final concentration of 0.001% (v/v). To measure the activity of *Es*LuxA, hexanal was added at the final concentration of 1% (v/v). After addition of 0.2 μM of *Pl*LuxAB or 6 μM of *Es*LuxA to the buffer, luminescence was measured using the Biotox-7BM luminometer.

### 2.6. Crystallization

Protein-containing fractions from SEC were pooled and, to prevent formation of disulfide bridges between surface-exposed cysteines, incubated with 15 mM iodoacetamide, which caused precipitation. Consequently, the protein was resuspended in a buffer containing 150 mM NaCl, 25 mM Tris-HCl and 15% glycerol, pH 8.0 and concentrated to 11.6 mg/ml for crystallization. Sitting drop vapor diffusion crystallization trials were set up using an NT8 robotic system (Formulatrix, USA) and stored at 20 °C. The drops contained 100 nL concentrated protein solution and 100 nL reservoir solution. The best crystals were obtained using the following solution as the precipitant: PEG3350 25%, lithium sulfate 0.2 M, BIS-TRIS 0.1 M pH 5.5. Cubic crystals appeared and reached the final size of ∼100 μm within a month. The crystals were harvested using micromounts and then flash-cooled and stored in liquid nitrogen.

### 2.7. Diffraction data collection and processing

The diffraction data were collected at 100 K at the European Synchrotron Radiation Facility (ESRF) beamline ID23-1 equipped with an EIGER2 S 16M detector (Dectris, Switzerland). Diffraction images were processed using XDS [42]. AIMLESS [43] was used to merge and scale the data. The data collection and processing statistics are reported in Table 2.

**Table 2.** Crystallographic data collection and refinement statistics.

|  |  |
| --- | --- |
| **Data collection** |  |
| Space group | P1 |
| Cell dimensions | - |
| $a, b, c$ (Å) | 51.8, 73.4, 94.0 |
| $\alpha, \beta, \gamma$ (°) | 88.7, 77.9, 87.1 |
| Wavelength (Å) | 0.9763 |
| Resolution (Å) | 91.89–2.71 (2.84–2.71) * |
| $R_{\text{merge}}$ (%) | 11.0 (31.0) * |
| $R_{\text{pim}}$ (%) | 11.0 (31.0) * |
| $\langle I/\sigma I \rangle$ | 4.0 (2.0) * |
| $CC_{1/2}$ (%) | 98.1 (89.4) * |
| Completeness (%) | 97.0 (96.2) * |
| Multiplicity | 1.8 (1.8) * |
| Unique reflections | 35,543 (4,681) * |
| **Refinement** |  |
| Resolution (Å) | 91.89–2.71 |
| No. reflections | 73,799 |
| $R_{\text{work}}/R_{\text{free}}$ (%) | 19.1/26.9 |
| No. atoms | - |
| Protein | 10897 |
| SO<sub>4</sub> ions | 20 |
| Water | 165 |
| Average $B$ factors (Å<sup>2</sup>) | - |
| Protein | 41.3 |
| SO<sub>4</sub> ions | 66.1 |
| Water | 26.4 |
| R.m.s. deviations | - |
| Protein bond lengths (Å) | 0.001 |
| Protein bond angles (°) | 0.33 |
| Ramachandran analysis | - |
| Favored (%) | 97.1 |
| Outliers (%) | 0.3 |

\* The data for the highest resolution shell is shown in parenthesis

### 2.8. Structure determination and refinement

The structure was solved using molecular replacement with MOLREP [44] and AlphaFold [45] model of *Es*LuxA as a search model. The resulting model was refined manually using Coot [46] and REFMAC5 [47].

### 2.9. Multi-angle light scattering (MALS)

SEC coupled with MALS (SEC-MALS) was done using a Superdex® 75 10/300 GL column (GE Healthcare Life Sciences, USA) and a combination of a UV-Vis Prostar 335 detector (Varian, Australia) and a miniDAWN detector (Wyatt Technology, USA) connected in this order. The *Es*LuxA sample (1 mg/ml in 120 μl) was loaded on the column equilibrated with filtered (0.1 μm) and degassed 25 mM Tris-HCl buffer, pH 8.0, containing 150 mM NaCl at 0.8 ml/min flow rate. Data were processed in ASTRA 8.0 (Wyatt Technology, USA) using dn/dc equal to 0.185 and protein extinction coefficient ε(0.1%) at 280 nm of 1.239.

### 2.10. Small-angle X-ray scattering (SAXS)

The SAXS experiments were carried out on the Rigaku instrument at the Moscow Institute of Physics and Technology (Dolgoprudny, Russia) as described previously [48,49]. SAXS profiles were obtained for the peak fraction after gel-filtration with protein concentration of 3.5 mg/ml, in a buffer containing 150 mM NaCl and 25 mM Tris-HCl, pH 8.0. For information on *Es*LuxA oligomerization polydispersity, we prepared models of the protein in monomeric, dimeric and tetrameric (dimer of dimers) states based on the crystal structure. The missing unstructured parts of the models were completed with fragments generated using AlphaFold [45]. Form-factors were calculated using CRYSOL [50]. The experimental data were approximated by superposition of form-factors using OLIGOMER [51] program from ATSAS software suite [52]. Details of SAXS measurements and data treatment are presented in Table S2. Guinier approximation, Kratky plot and pair distribution function are shown in Fig. S1.

## 3. Results

### 3.1. Phylogenetic analysis of luxA-related genes

We began our study with reanalysis of the available genomic data. We obtained the sequences of the monooxygenase fold proteins from InterPro and constructed a phylogenetic tree (Fig. 1). In accordance with the findings by Vannier and colleagues [30], we observed that *luxA* and *luxB* genes form two well-defined clusters, separate from other monooxygenase family members. *luxF* is highly dissimilar from *luxA* and *luxB* and is unlikely to be a direct descendant of a luciferase gene (Fig. 1). Several *luxA*-like genes are observed near the branching point of *luxA* and *luxB* clusters as parts of diverse operons also including *luxC*-like, *luxD*-like and *luxE*-like genes (Fig. 1). Compared to the findings of Vannier and colleagues [30], the genes from *Leptospira fletcheri, Leptospira fluminis, Leptospira perolatii, Nocardia brasiliensis* and *Nocardia uniformis* appear to be new (Table S3).

**Fig. 1.**
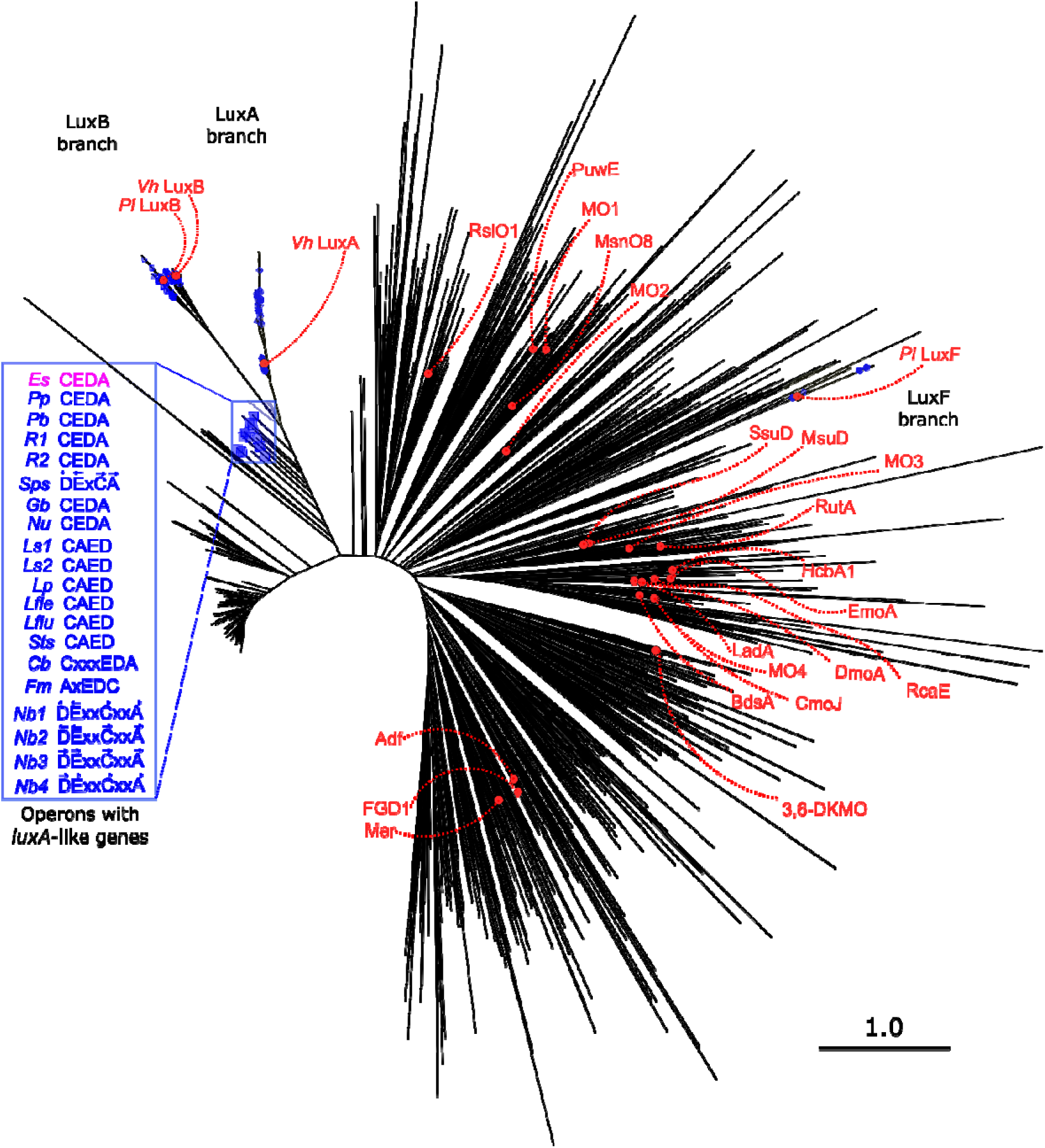
Phylogenetic tree of *luxA, luxB* and *luxA*-like genes. The genes with *luxC*/*luxD*/*luxE* in vicinity are marked in blue. The inset shows the gene order in the putative luciferase operons; details are presented in Table S3. Proteins with available experimentally determined structures are labeled in red; details are presented in Table S4.

### 3.2. Luminescence properties of E. salina LuxA in vivo and in vitro

Willing to test the properties of the proteins encoded by *luxA*-like genes experimentally, we synthesized a gene encoding putative *E. salina* luciferase LuxA (*Es*LuxA, GenBank ID WP_106392368, sequence identity to *V. harveyi* LuxA is 41%) with a C-terminal histidine tag and cloned it into the pET11 plasmid. *E. coli* cells, transformed with the plasmid, displayed moderate luminescence upon addition of various aldehydes to the media (Fig. 2). Compared to canonical heterodimeric luciferases from *A. fischeri* (*Af*LuxAB) and *P. luminescens* (*Pl*LuxAB), which are most active with longer-chain aldehydes, *Es*LuxA was most active with medium-chain C4-C9 aldehydes. Normalized for the aldehyde concentration and cell density, the activity of *Es*LuxA was around 10^-5^ of that of *Af*LuxAB and *Pl*LuxAB.

**Fig. 2.**
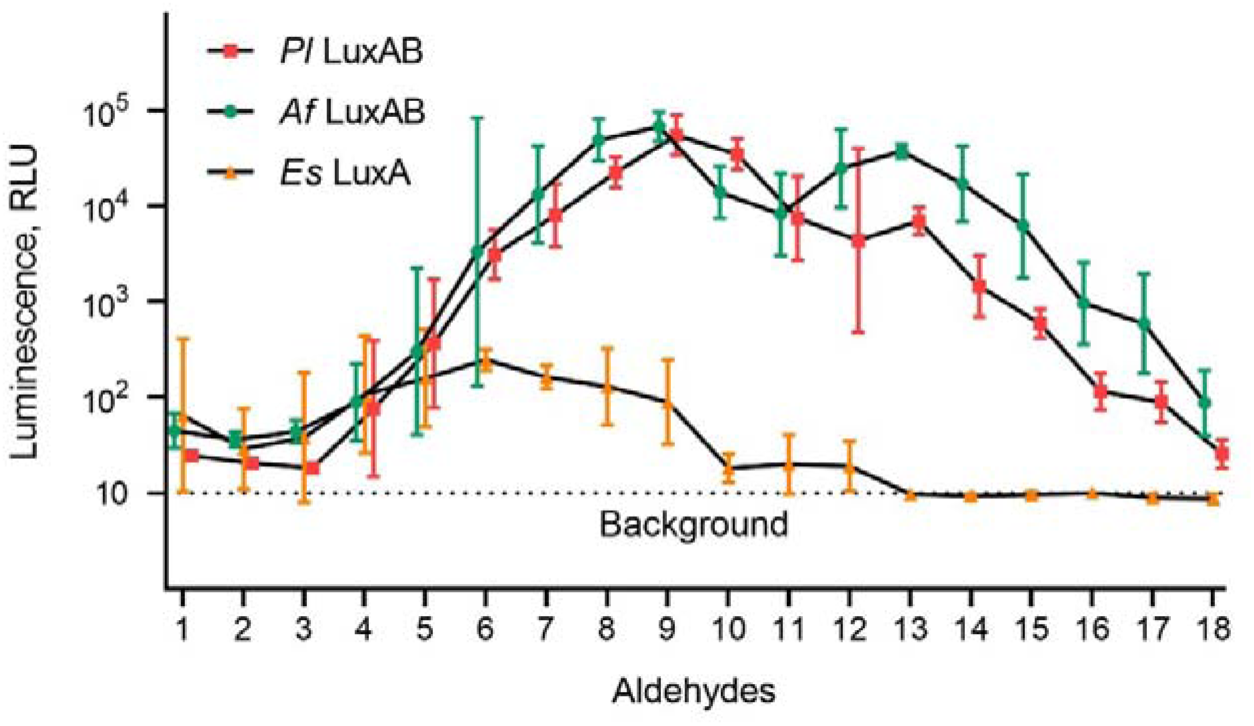
Dependence of luminescence on the aldehyde aliphatic chain length for *Pl*LuxAB (*E. coli* MG1655 pABX), *Af*LuxAB (*E. coli* MG1655 pF2), and *Es*LuxA (*E. coli* BL21 (DE3) pSluxA). Aldehydes were used at the final concentration of 1 mM. Cells with *Es*LuxA were assayed at 40× optical density of cells with *Pl*LuxAB or *Af*LuxAB because of their low luminescence. Data points and error bars correspond to means and standard deviations of logarithms of luminescence values determined in three independent experiments. RLU is a relative luminescence unit.

Next, we tested the dependence of luminescence on the aldehyde concentration *in vivo*. Apparent Michaelis constants (*K*_M_) determined for *Es*LuxA-expressing cells are 9±3 mM both for hexanal and nonanal, whereas for *Pl*LuxAB-expressing cells *K*_M_ is much lower for nonanal (25±2 µM) than for hexanal (7±2 mM).

Finally, we tested luciferase activity of purified *Es*LuxA and *Pl*LuxAB *in vitro* (Figure 3) using the optimal substrates identified in *in vivo* measurements: hexanal for *Es*LuxA and decanal for *Pl*LuxAB. The maximum level of luminescence for *Es*LuxA was ∼2 RLU above the background, while for *Pl*LuxAB it was ∼10 000 RLU. Taking into account enzyme concentrations of 300 µg/ml and 20 µg/ml for *Es*LuxA and *Pl*LuxAB, respectively, the level of specific luminescence of *Es*LuxA is approximately 10^-5^ of that of *Pl*LuxAB, in agreement with data from *in vivo* experiments.

**Fig. 3.**
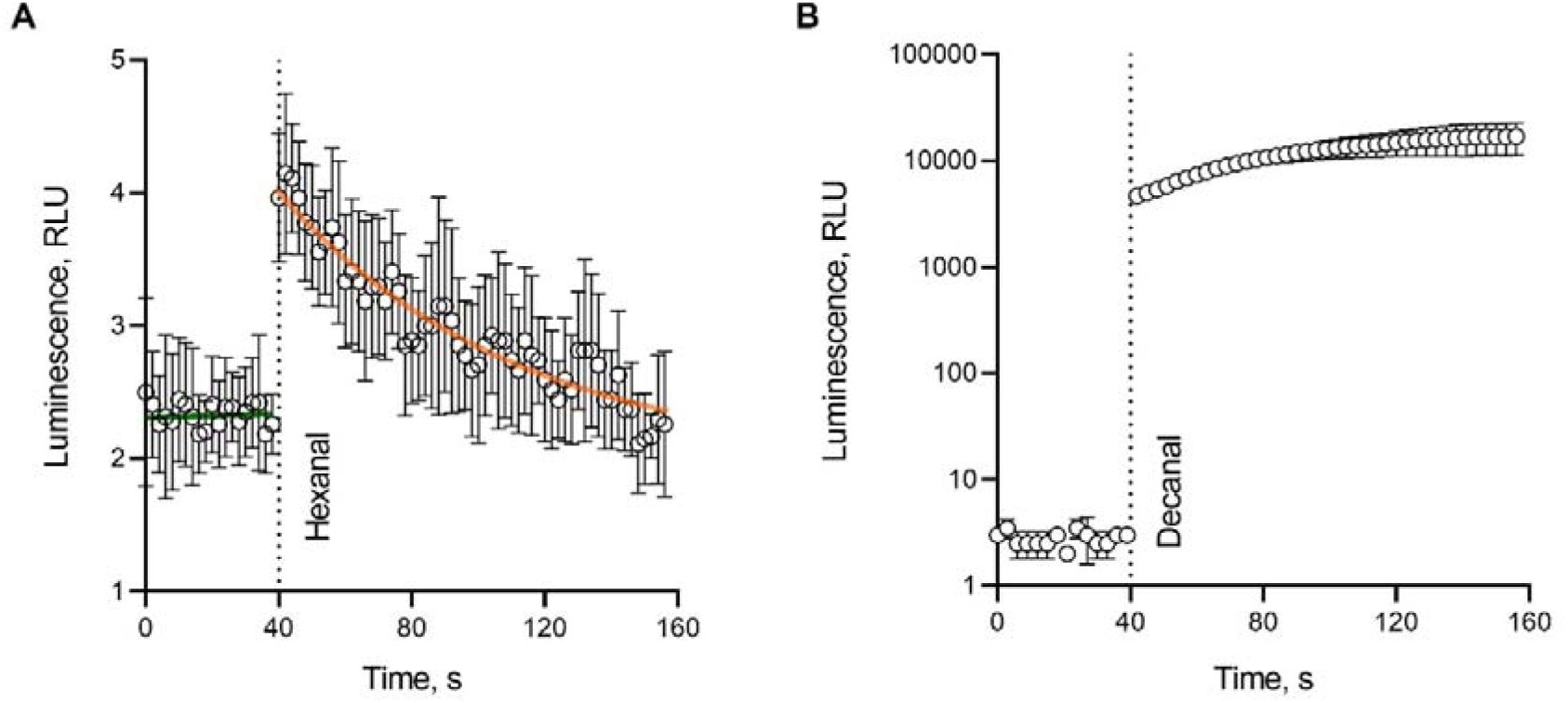
Luminescence kinetics for luciferases *Es*LuxA (A) and *Pl*LuxAB (B) *in vitro*. Aldehydes were added to the samples at 40 s. Green and orange lines in panel A correspond to fitted approximations of data points before and after addition of hexanal, correspondingly. Because of significant difference in luminescence intensity of luciferases EsLuxA и PlLuxAB, luminescence scale is linear in panel A and logarithmic in panel B. Data points and error bars correspond to means and standard deviations of luminescence values obtained in 9 (A) and 3 (B) independent measurements. RLU is a relative luminescence unit.

### 3.3. Structural characterization of Enhygromyxa salina LuxA

To gain insight into the *Es*LuxA structural properties, we overexpressed the protein in *E. coli* and purified it using metal-affinity and size-exclusion chromatography. The purified protein was colorless and, presumably, in the apo-form. We determined the crystal structure of *Es*LuxA at the resolution of 2.71 Å. The P1 unit cell contains two *Es*LuxA homodimers. *Es*LuxA has an (α/β)_8_ barrel fold characteristic for LuxA/LuxB and other monooxygenase family members [17] (Fig. 4). The structures of all *Es*LuxA protomers within the crystal are essentially identical with the exception of the amino acids 264-293, corresponding to the mobile loop of the α-subunit of classical heterodimeric bacterial luciferases (residues 259-283 in *Vh*LuxA, Fig. 4A and B). Presence of this elongated partially unstructured loop confirms annotation of the *E. salina luxA* gene product as LuxA-like, rather than LuxB-like. While the loop is disordered in protomers B and D, some of its residues form α-helices in protomers A and C (Fig. 4B). No putative cofactors or substrates are observed in the electron densities; a single sulfate group is observed in the *Es*LuxA structure at the place of the FMNH_2_’s phosphate in the *Vh*LuxAB structure.

**Fig. 4.**
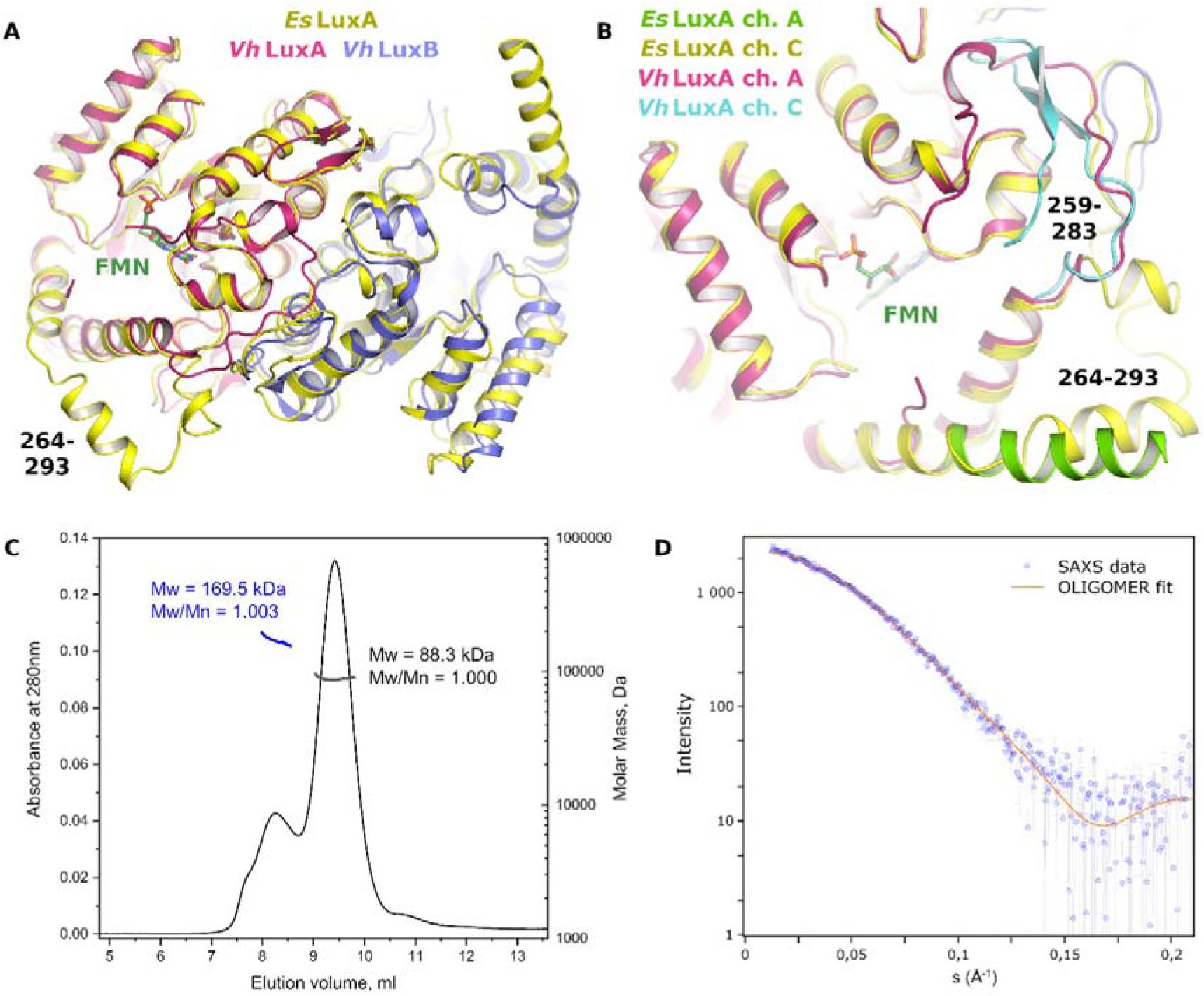
Structural properties of *Es*LuxA in crystal and in solution. (A) Overlay of the crystallographic structure of homodimeric *Es*LuxA with heterodimeric *Vh*LuxAB (PDB ID 3FGC, [17]). (B) Overlay of the active site of *Es*LuxA with that of *Vh*LuxAB. For both complexes, different molecules in the crystallographic unit cells (chains A and C) show different arrangements of the elongated loop (α-helix and coil in *Es*LuxA, β-hairpin and coil or just coil in *Vh*LuxA). (C) SEC-MALS chromatogram. Two major fractions correspond to *Es*LuxA dimers and tetramers. (D) SAXS data (blue) and theoretical scattering curve (orange) corresponding to a mixture of 82% dimers and 18% tetramers of *Es*LuxA.

To determine the oligomeric state of *Es*LuxA in solution, we conducted SEC-MALS and small-angle X-ray scattering (SAXS) experiments. While the MW of His-tagged *Es*LuxA monomer is expected to be 42.2 kDa, SEC-MALS revealed that the major fraction contained the species with MW of 88.3 kDa, and the minor fraction contained the species with MW of 169.5 kDa (Fig. 4C), corresponding to *Es*LuxA dimers and tetramers, respectively. In accordance with this, the SAXS profile could be reasonably fitted with the theoretical scattering of 82% to 18% mixture of *Es*LuxA dimers and tetramers, respectively (Fig. 3D), which is in accordance with the MALS data.

## 4. Discussion

Recently, Vannier and colleagues identified *luxA*-like genes in species that were not reported to display luminescence [30]. Phylogenetic analysis shows that previously known *luxA* and *luxB* probably arose from duplication of some protoluciferase gene and underwent asymmetric divergence [53]: all *luxB* have approximately the same, relatively large, distance to the branching point, whereas some of the *luxA* genes are relatively close to it (Fig. 1). *luxA*-like genes are similarly closest to the *luxA*/*luxB* branching point. Given that *Es*LuxA and related proteins all have an elongated ‘mobile loop’ covering the active site, characteristic for α, but not β subunits of heterodimeric luciferases [30], it is reasonable to denote them *luxA* and not *luxB*. At the same time, it is important to keep in mind that the main function of these genes and proteins might be not luminescence *per se*, and thus they might represent protoluciferases rather than actual luciferases.

Here, we showed that the *luxA*-like gene from *E. salina* indeed encodes a functional homodimeric luciferase, *Es*LuxA, which, in contrast with *Vh*LuxA, remains monomeric [22]. Crystallographic structure of *Es*LuxA is similar to those of *Vh*LuxAB heterodimer and *Vh*LuxB homodimer. MALS and SAXS analyses show that in solution *Es*LuxA preferentially forms dimers, with a minor addition of tetramers.

*E. salina* is a slightly halophilic myxobacterium isolated from coastal areas of Japan [54] that has not been described as being luminescent. Accordingly, we found that *Es*LuxA is dim; its level of luminescence is several orders of magnitude lower compared to classical heterodimeric luciferases (Fig. 2 and 3). Coexpression of *E. salina luxA* with *P. luminescens luxB* or *A. logei luxB* did not lead to an increase in luminescence (data not shown). Previously reported attempts to hybridize other bacterial luciferases were also not successful [55]. Evidently, the subunit interaction interface is important for proper functioning of bacterial luciferases [56] and evolved for preferential binding of the native partner.

The optimal aldehyde tail length for *Es*LuxA is around 6 carbon atoms (Fig. 2); in contrast, heterodimeric luciferases *Af*LuxAB and *Pl*LuxAB are more active with aldehydes with chain lengths of 8-10 carbons (Fig. 2 and refs. [55,57]). Due to low activity of *Es*LuxA, it was difficult to assess the properties of the purified enzyme. Previously, the activity and *K*_M_ values of the *A. fisheri* luciferase obtained *in vivo* and *in vitro* were found to be almost the same for aldehydes shorter than octanal, while long chain aldehydes resulted in higher *K*_M_ values *in vivo* [58]. We found that for *Pl*LuxAB-expressing cells the *K*_M_ constant is ∼300 times lower for nonanal (25±2 µM) than for hexanal (7±2 mM). Interestingly, for *Es*LuxA-expressing cells, there is no difference between hexanal and nonanal, as the *K*_M_ constants for both are 9±3 mM.

Altogether, our results show that *Es*LuxA is dim and rather nonselective, and thus is an unusual luciferase. Possible explanations for these findings are a) accidental loss of the *luxB* gene in the *lux*-operon of *E. salina*; b) requirement of native *luxC*, *luxD* and *luxE* genes for optimal functioning, and/or preference for an unorthodox aldehyde as the preferred substrate; c) that the main function of *Es*LuxA is not to emit light, but rather to be a terminal oxidase, providing an alternate respiratory pathway for cells grown under conditions where the cytochrome system is functionally impaired [59].

To summarize, our work paves the way for discovery of new unusual bacterial luciferases and for description of new luminescent microorganisms. Homodimeric luciferase encoded by a single gene is also promising for development of new tools for molecular biology: bacterial luciferases are used ubiquitously to construct whole cell biosensors or molecular reporters [60–62], yet their heterodimeric nature complicates heterologous expression and engineering efforts [28,29,63].

## Supporting information

Supporting text, supporting tables 1-4 and supporting figure 1 are available.

## Data availability

Generated data are made available either as supporting information or via public databases. Atomic coordinates and structure factors for the reported crystal structure have been deposited in the Protein Data Bank [64] under the accession code 8CBB. Small angle scattering data have been deposited in the SASBDB [65] under the accession code SASDQA7.

## Acknowledgements

The diffraction experiments were performed on the beamline ID23-1 at the European Synchrotron Radiation Facility (ESRF), Grenoble, France [66]. We are grateful to Sergey Bukhdruker for conducting preliminary diffraction tests and to the ID23-1 beamline staff of ESRF for providing assistance in using the beamline.

