## Supporting Information for "*luxA* gene from *Enhygromyxa salina* encodes a functional homodimeric luciferase"

Supporting Text: Detailed description of cloning for pSluxA, pABX, pABX-T7, and pLuxG-T7 plasmids; nucleotide sequence optimized for *Es*LuxA expression in *E. coli.*

Supporting Tables 1-4

Supporting Figure 1

**Detailed description of cloning for pSluxA, pABX, pABX-T7, and pLuxG-T7 plasmids**

The gene encoding the putative LuxA protein from *E. salina* (*Es*LuxA, GenBank ID WP_106392368) with C-terminal 6×His-tag was optimized for *E. coli* expression and synthesized *de novo*; the nucleotide sequence is presented below. The gene was cloned into the pET11 expression vector (Novagen, Merck, Germany) using BamHI and NdeI restriction sites, the resulting plasmid was named pSluxA.

The pABX plasmid harboring the *luxAB* genes from *P. luminescens* (GenBank ID AF403784.1) under control of $P_{lac}$ was prepared by linearizing the pUC19 vector by KpnI restriction endonuclease, amplifying the *luxAB* fragment from the pXen7 plasmid [1], and ligating the resulting fragments with the use of Gibson assembly [2].

To construct pABX-T7, the *luxAB* fragment from *P. luminescens* was amplified from pXen7 (six codons of His added by primer), then it was cloned into the pET15b vector at the BamHI/NcoI sites (restriction sites in PCR-fragment were added by the primers, underlined in Table S1). As a result, in pABX-T7, the *luxAB* genes encoding LuxA with a 6×His tag at the N-terminus and LuxB are located under the T7 promoter.

pLuxG-T7 plasmid was prepared by linearizing the pET15b vector by NdeI restriction endonuclease, amplifying the *luxG* fragment from the chromosomal DNA of *V. aquamarines* (GenBank ID PP639076), and ligating the resulting fragments using Gibson Assembly.

Oligonucleotides used for plasmid preparation may be found in Table S1.

### Nucleotide sequence optimized for *Es*LuxA expression in *E. coli*

ATGACCCAGAGCGAGGACATTAAATGGGGTATGTTTCTGAATACCGCACGTCCGCCTCAGTT
TACCGAACGTCGTGTTCTGGAAAATGCAAAATTCTATGGTCAGGTTGCAGAAGAAATGGGTT
TTGAAAGCGCATGGATGCTGGAACATCATTTTACCGATTATGGTCTGTGTGGTAGCCCGATG
GTTATGGCAAGCTATATTCTGGGTGCAACCCGTCGTATTAAAGTTGGCACCGCAATTAACAT
TCTGCCGCTGGAACACCCGGTTCGTCTGGCAGAACAGGCAGCACTGCTGGATCAGCTGAGTG
ATGGTCGTTTTATTCTTGGTATTGGTCGCGGTTTCTTCGATAAAGATTTTACCGTTTTTGGC
GTGGACATCCATGATACCCGTGCACTGACCCATAACTATTATGACATTATGCAAGAGGCATG
GACCAAAGGTGTTGTTGGTTCAGATGGTCCGTTTCTGAACTTTCCGCCTGTTCCGGTTAATC
CGCGTCCGTATAGCGATAAAATGCCGATGGTTTGTGCAGCAATGAGCCCGAGCACCATTGAA
TGGGCAGCAAAAAATGGTCTGCCGATGATTATGCAGCATGACATTGAACATAATGAGAAAGC
CAGCAACGTTGAACTGTATCGTGCACTGGCCGAAGAACATGGTCACGATCCTGATGGTATCG
AACATACCATTGCCATGATTGTTGCAGTTGATCCTGATCGTGAACGTGTTCGTGAAGAATGT
CGTCATTATCTGAACTGGTTTGAAGATGCAGTTGAAAAAGCCCAGAACATTATTGATATTGT
GCGTGAACATGGCGTGGAATGTTATGATTGGCATCTGCGTAAATGGCGTGAAGCAGTTATTA
AAGGTGATACCGCCATTAGCAAAGTGGTTGATAATCTGCTGCGTCTGAATGCCATTGGTACA
CCGGAAGATGCCATTGAAACCATTCAGCATGTTATTGATGTGACGGGTGTTAAACGTGTTGT
GGTTGGTTTTGAAGCGATTGGTGATCGTGATCGCGTGCTGGAAAGCATGAAACTGTTTGATG
AACAGGTTCGTCCGCATATTCGTGGTGCAAAAATTGGTAGCCATCACCATCATCATCATTAA

Nucleotides corresponding to C-terminal GSHHHHHH tag are underlined.

**Table S1.** Primers used in the study. Nucleotides corresponding to restriction sites are underlined.

| **Name** | **Sequence** | **Note** |
|---|---|---|
| xenLuxBR-pUC | CCAGTGAATTCGAGCTCGGTACCGTTGCTGCCATATTCTTTTA | For pABX construction |
| xenLuxAD-pUC | AGAGGATCCCCGGGTACATGAAATTTGGAAACTTTTTGCTTA | |
| luxAD | G<u>CCATGG</u>GCCATCATCATCATCATCACAGCGGCAGCGGCATGAAATTTGGAAACTTTTTGCTTAC | For pABX-T7 construction |
| luxBR | GTT<u>GGATCC</u>ATATTCTTTTACTACATGTGGTACT | |
| luxGD | CCGCGCGGCAGCCATATGTTATGTACGGTAGAAAAAATAGAACC | For pLuxG-T7 construction |
| luxGR | AGCCGGATCCTCGAGCAGTTATAGGTAAGCGAATGCGTCAGC | |

**Table S2.** SAXS experimental details and data evaluation summary.

| | |
|---|---|
| ***(a)* Sample details** | |
| Description of sequence | Homodimeric luciferase *Es*LuxA<br>GenBank ID WP_106392368 with C-terminal 6×His tag (host organism *Enhygromyxa salina*) |
| Chemical formula [1] | $C_{1879}H_{2904}N_{530}O_{545}S_{18}$ |
| Molecular mass (kDa) [1] | 42216.03 |
| Approximate volume (Å$^3$) [1] | 51079 |
| Partial specific volume ν (cm$^3$ g$^{-1}$) [1] | 0.7286 |
| Mean solute and solvent SLD (10$^{-6}$ Å$^{-2}$) [1] | 9.40; 12.48 |
| Mean scattering contrast Δρ (10$^{-6}$ Å$^{-2}$) [1] | 3.08 |
| Extinction coefficient $\varepsilon_{280nm}$ (M$^{-1}$ cm$^{-1}$) | 51910 |
| Sample concentration (mg ml$^{-1}$) | 3.2 |
| Solvent composition | 150 mM NaCl, 25 mM Tris-HCl (pH 8.0) |
| **(*b*) SAS data collection parameters** | |
| Instrument | Rigaku MicroMax-007HF (MIPT, Dolgoprudny, Russia) |
| Wavelength (Å) | 1.5406 |
| Beam geometry (FWHM diameter, sample-to-detector distance) | 0.3 mm, 2.0 m |
| Sample configuration | Glass capillaries with diameters of 1.2 mm and 1.3 mm for protein sample and buffer, correspondingly |
| *q*-measurement range (Å$^{-1}$) | 0.007 – 0.215 |
| *q*-scaling method | Calibration standard: silver behenate powder |
| Basis for normalization to constant counts | To path length and transmission (measured by direct beam counter) |
| Exposure time | 140 min |
| Sample temperature (°C) | 20 |
| **(*c*) Software employed for SAS data reduction, analysis and interpretation** | |
| SAS data subtraction and Guinier analysis | PRIMUSqt from ATSAS 2.8.4 |
| Calculation of ε from sequence | ProtParam: https://web.expasy.org/protparam/ |
| Calculation ν values from chemical composition | Peptide Property Calculator: http://biotools.nubic.northwestern.edu/proteincalc.html |
| Calculation of scattering length density (SLD) values from chemical composition | SLD calculator: http://www.ncnr.nist.gov/resources/activation/ |
| *P*(*r*) calculation | GNOM 5.0 from ATSAS 2.8.4 |
| Atomic structure modelling and fitting the data | AlphaFold 2; CRYSOL; OLIGOMER |
| **(*d*) Structural parameters** | |
| Guinier analysis | |
| *I*(0) (a.u.) | 0.249 ± 0.002 |
| $R_g$ (Å) | 32.9 ± 0.4 |
| *q*-range (Å$^{-1}$) ($qR_g$ range) | 0.0125 – 0.0395 (0.41 – 1.30) |
| *P*(*r*) analysis | |
| *I*(0) (a.u.) | 0.249 ± 0.002 |
| $R_g$ (Å) | 33.4 ± 0.5 |
| $d_{max}$ (Å) | 125 |
| *q*-range (Å$^{-1}$) | 0.0125 – 0.210 |
| $q_{min}\, d_{max} / \pi$ | 0.497 |
| Total quality estimate (GNOM) | 0.656 |

| | |
|---|---|
| Volume ($V_P$) ($nm^3$) | 137.7 (± 10%) |
| MW calculated from $V_P$ (kDa) | 113.8 (± 10%) |
| Experimental MW (kDa) | 99 ± 4 (in accordance with weights from OLIGOMER fit, see section *(e)*) |
| ***(e)* Atomistic modelling** | |
| Method | OLIGOMER program from ATSAS was used to approximate experimental data by a set of curves calculated for monomers, dimers and tetramers (dimers of dimers) of *Es*LuxA using CRYSOL. The models of *Es*LuxA monomers, dimers and tetramers were built based on the crystal structure. The residues missing in the crystal structure were added using AlphaFold v.2. |
| $q$-range for fitting | 0.0125 – 0.210 |
| Contrast of the solvation shell ($10^{-6}$ Å$^{-2}$); Average atomic radius (Å) | 0.844; 1.616 |
| Volume fractions | Monomers: 0.005 ± 0.031<br>Dimers: 0.818 ± 0.039<br>Dimers of dimers: 0.177 ± 0.015 |
| $\chi^2$ value | 1.00 |
| ***(f)* Data and model deposition IDs** | |
| | Homodimeric luciferase *Es*LuxA<br>SASBDB SASDQA7 |

[1] These values were calculated from the protein sequence without taking into account possible ligands.

**Table S3.** Information about the *luxA*-like genes.

| **Abbreviation** | **Organism** | **GenBank accession ID** | **Gene order** | **Lineage** |
|---|---|---|---|---|
| *Es* | *Enhygromyxa salina* | WP_106392368.1 | CEDA | *Bacteria; Proteobacteria; Deltaproteobacteria; Myxococcales; Enhygromyxa* |
| *Pp* | *Plesiocystis pacifica* | WP_006970588.1 | CEDA | *Bacteria; Proteobacteria; Deltaproteobacteria; Myxococcales; Nannocystineae; Nannocystaceae; Plesiocystis* |
| *Pb* | *Pseudomonadales bacterium* | MAA60901.1 | CEDA | *Bacteria; Proteobacteria; Gammaproteobacteria; Pseudomonadales* |
| *R1* | *Rhizobacter sp. Root1221* | KQV90440.1 | CEDA | *Bacteria; Proteobacteria; Betaproteobacteria; Burkholderiales;Rhizobacter* |
| *R2* | *Rhizobacter sp. Root1221* | WP_082568857.1 | CEDA | *Bacteria; Proteobacteria; Betaproteobacteria; Burkholderiales; Rhizobacter* |
| *Sps* | *Spongiibacter sp. KMU-166* | WP_168452140.1 | D<E<xCA | *Bacteria; Proteobacteria; Gammaproteobacteria; Cellvibrionales; Spongiibacteraceae; Spongiibacter* |
| *Gb* | *Gammaproteobacteria bacterium 45_16_T64* | OUS24261.1 | CEDA | *Bacteria; Proteobacteria; Gammaproteobacteria* |
| *Nu* | *Nocardia uniformis* | WP_170264378.1 | CEDA | *Bacteria; Actinobacteria; Corynebacteriales; Nocardiaceae; Nocardia* |
| *Ls1* | *Leptospira santarosai* | WP_004465116.1 | CAED | *Bacteria; Spirochaetes; Leptospirales; Leptospiraceae; Leptospira* |
| *Ls2* | *Leptospira santarosai* | WP_004490151.1 | CAED | *Bacteria; Spirochaetes; Leptospirales; Leptospiraceae; Leptospira* |
| *Lp* | *Leptospira perolatii* | WP_100713053.1 | CAED | *Bacteria; Spirochaetes; Leptospirales; Leptospiraceae; Leptospira* |
| *Lfle* | *Leptospira fletcheri* | WP_135769088.1 | CAED | *Bacteria; Spirochaetes; Leptospirales; Leptospiraceae; Leptospira* |
| *Lflu* | *Leptospira fluminis* | WP_135814401.1 | CAED | *Bacteria; Spirochaetes; Leptospirales; Leptospiraceae; Leptospira* |
| *Sts* | *Streptomyces sp. SLBN-118* | WP_142212242.1 | CAED | *Bacteria; Actinobacteria; Streptomycetales; Streptomycetaceae; Streptomyces* |
| *Cb* | *Chloroflexi bacterium* | TMC11770.1 | CxxxEDA | *Bacteria; Chloroflexi* |
| *Fm* | *Candidatus Frankia meridionalis* | WP_165486026.1 | AxEDC | *Bacteria; Actinobacteria; Frankiales; Frankiaceae; Frankia* |
| *Nb1* | *Nocardia brasiliensis* | WP_042263712.1 | D<E<xxCxxA | *Bacteria; Actinobacteria; Corynebacteriales; Nocardiaceae; Nocardia* |
| *Nb2* | *Nocardia brasiliensis* | WP_195134718.1 | D<E<xxCxxA | *Bacteria; Actinobacteria; Corynebacteriales; Nocardiaceae; Nocardia* |
| *Nb3* | *Nocardia brasiliensis* | WP_014985126.1 | DExxCxxA | *Bacteria; Actinobacteria; Corynebacteriales; Nocardiaceae; Nocardia* |
| *Nb4* | *Nocardia brasiliensis* | WP_029901948.1 | DExxCxxA | *Bacteria; Actinobacteria; Corynebacteriales; Nocardiaceae; Nocardia* |

**Table S4.** Representative luciferase-like monooxygenase family members with available experimentally determined structures.

| Abbreviation | Description | Organism | Primary publication | PDB ID |
|---|---|---|---|---|
| *Vh* LuxA | Luciferase subunit α | *Vibrio harveyi* | [3] | 3FGC_A |
| *Vh* LuxB | Luciferase subunit β | *Vibrio harveyi* | [3] | 3FGC_B |
| *Pl* LuxB | Luciferase subunit β | *Photobacterium leiognathi* | no publication identified | 6FRI_A |
| *Pl* LuxF | Non-fluorescent flavoprotein FP390 LuxF | *Photobacterium leiognathi subsp. leiognathi* | [4] | 1NFP_A |
| 3,6-DKMO | 3,6-diketocamphane monooxygenase | *Pseudomonas putida* | [5] | 4UWM_A |
| Adf | Alcohol dehydrogenase | *Methanoculleus thermophilus* | [6] | 1RHC_A |
| BdsA | Dibenzothiophene sulfone monooxygenase | *Bacillus subtilis* | [7] | 5TLC_A |
| CmoJ | Putative monooxygenase | *Bacillus subtilis subsp. subtilis str. 168* | no publication identified | 6ASK_A |
| DmoA | Dimethylsulfide monooxygenase | *Hyphomicrobium sulfonivorans* | [8] | 6AK1_A |
| EmoA | EDTA monooxygenase | EDTA-degrading bacterium *BNC1* | [9] | 5DQP_A |
| FGD1 | Glucose-6-phosphate dehydrogenase | *Rhodococcus jostii RHA1* | [10] | 5LXE_A |
| HcbA1 | Hexachlorobenzene monooxygenase | *Nocardioides sp. PD653* | [11] | 6LR1_A |
| LadA | Long-chain alkane monooxygenase | *Geobacillus thermodenitrificans* | [12] | 3B9N_A |
| Mer | 5,10-methylenetetrahydromethanopterin reductase | *Methanothermobacter thermautotrophicus* | [13] | 1F07_A |
| MO1 | Monooxygenase | *Agrobacterium tumefaciens* | no publication identified | 2I7G_A |
| MO2 | Luciferase-like monooxygenase | *Bacillus cereus* | no publication identified | 2B81_A |
| MO3 | Putative Luciferase-like Monooxygenase | *Bacillus cereus ATCC 10987* | no publication identified | 3RAO_A |
| MO4 | Nitrilotriacetate monooxygenase | *Burkholderia pseudomallei 1710b* | no publication identified | 3SDO_A |
| MsnO8 | Luciferase-like monooxygenase | *Streptomyces bottropensis* | [14] | 4US5_A |
| MsuD | Alkanesulfonate monooxygenase | *Pseudomonas fluorescens Pf0-1* | [15] | 7JV3_A |
| PuwE | Monooxygenase (OX) domain of hybrid polyketide/non-ribosomal peptide synthetase | *Cylindrospermum alatosporum CCALA 988* | no publication identified | 6KET_A |
| RcaE | Riboflavin lyase | *Herbiconiux* | no publication identified | 5W48_A |
| RslO1 | Luciferase-like monooxygenase | *Streptomyces bottropensis* | [16] | 7BIP_A |
| RutA | Pyrimidine monooxygenase | *Escherichia coli* | [17] | 5WAN_A |
| SsuD | Alkanesulfonate monooxygenase | *Escherichia coli* | [18] | 1M41_A |

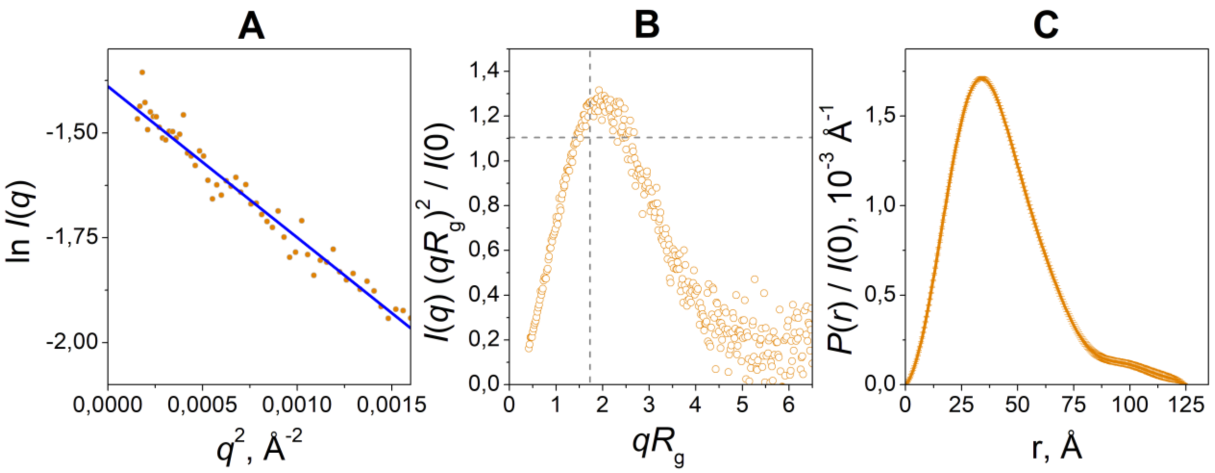


**Fig. S1.** Analysis of SAXS data for *Es*LuxA. (A) Guinier approximation. (B) Dimensionless Kratky plot. Dashed lines correspond to $qR_g = \sqrt{3}$ and to $(qR_g)^2\ I(q)\ /\ I(0) = 1.104$. (C) Pair distance distribution function $P(r)$.
